# WEE1 kinase is a therapeutic vulnerability in CIC-DUX4 undifferentiated sarcoma

**DOI:** 10.1101/2021.06.21.448439

**Authors:** Rovingaile Kriska Ponce, Nicholas J. Thomas, Nam Q. Bui, Tadashi Kondo, Ross A. Okimoto

## Abstract

CIC-DUX4 rearrangements define an aggressive and chemotherapy-insensitive subset of undifferentiated sarcomas. The CIC-DUX4 fusion drives oncogenesis through direct transcriptional upregulation of cell cycle and DNA replication genes. Notably, CIC-DUX4- mediated *CCNE1* upregulation compromises the G1/S transition, conferring a potential survival dependence on the G2/M cell cycle checkpoint. Through an integrative transcriptional and kinase activity screen using patient-derived specimens, we now show that CIC-DUX4 sarcomas depend on the G2/M checkpoint regulator, WEE1, as an adaptive survival mechanism. Specifically, CIC-DUX4 sarcomas depend on WEE1 activity to limit DNA damage and unscheduled mitotic entry. Consequently, genetic or pharmacologic WEE1 inhibition *in vitro* and *in vivo* leads to rapid DNA damage-associated apoptotic induction of patient-derived CIC-DUX4 sarcomas. Thus, we identify WEE1 as an actionable therapeutic vulnerability in CIC-DUX4 sarcomas.

## Introduction

Chromosomal rearrangements that create transcription factor (TF) fusion oncoproteins are attractive cancer-specific therapeutic targets (1). For example, mechanistic studies that provide insight into fusion oncoprotein stability have led to the development of precision-based therapies that directly degrade these driver oncoproteins (1–3). Beyond direct degradation, an alternative strategy to overcome TF fusion dependence in cancer is through the identification of key transcriptional targets that TF fusions directly control to drive malignant progression. To this end, we have recently identified a molecular dependence on the CCNE/CDK2 complex in CIC-DUX4 sarcoma, whereby CIC-DUX4 transcriptionally upregulates *CCNE1* to drive sarcoma growth and survival (4). Therapeutic intervention with CDK2 inhibitors leads to apoptotic induction in patient-derived CIC-DUX4 models (4). These findings provided a mechanism-based therapeutic strategy to limit CIC-DUX4 sarcoma progression, which remains an aggressive and lethal disease.

In other cancers, increased expression of *CCNE1* through transcriptional upregulation or amplification often leads to a deficient G1/S checkpoint, thus enhancing DNA replication stress and genomic instability (5–11). Cancer cells adapt to these high replication stress states through an increased dependence on cell replication checkpoints that enable accurate DNA repair, scheduled mitotic entry, and survival (8, 12). When these key checkpoints are compromised, cancer cells undergo replication stress driven mitotic catastrophe and death (13, 14). One critical cell cycle checkpoint regulator is the WEE1 kinase, which modulates CDK1 and CDK2 activity through direct inhibitory phosphorylation (15, 16). Functionally, phosphorylation at Tyrosine 15 (Y15) on CDK1 by WEE1 delays progression at the G2/M checkpoint (15, 17), restricting premature entry into mitosis. Thus, as an adaptive mechanism to CCNE1-associated replication stress, WEE1 can enhance tumor cell survival. Through an integrative transcriptional and functional approach, we investigated how CIC-DUX4 sarcomas survive in a CCNE1- mediated high replication stress state through increased dependence on the WEE1 kinase. Moreover, we demonstrate that WEE1 is a therapeutic vulnerability in CIC-DUX4 sarcomas that can be targeted with clinically advanced WEE1 inhibitors.

## Results/Discussion

Prior studies have identified cell cycle regulators as direct transcriptional targets of the CIC-DUX4 fusion oncoprotein (4, 18, 19). One key CIC-DUX4 transcriptional target is *CCNE1,* which regulates the G1/S cell cycle transition (4, 20). CIC-DUX4-dependent *CCNE1* transcriptional upregulation compromises the G1/S checkpoint and confers a molecular and therapeutic dependence on the CCNE1/CDK2 complex (4). In order to identify additional actionable therapeutic targets in CIC-DUX4 sarcomas we integrated CIC-DUX4-dependent gene expression changes with functional kinase activity screens in CIC-DUX4 patient-derived cells to nominate candidate kinases that enable CIC-DUX4 survival (Figure 1A). Specifically, we performed a comparative transcriptional analysis using a validated dataset (GSE60740) comprised of CIC-DUX4 patient-derived cells (IB120) with or without genetic silencing of CIC-DUX4 (19). Through a previously described approach (4) we identified 165 putative CIC-DUX4 responsive genes, 43 of which contained the highly conserved CIC-binding motif (T[G/C]AATG[A/G]A) within −2 kb and +150 bp of the transcription start site (21). This analysis enabled us to identify high confidence CIC-DUX4-dependent gene expression changes in endogenous expressing CIC-DUX4 cells. In concordance with prior findings, we noted significant enrichment in genes that regulate the cell cycle, including G1/S transition, mitosis, and cell cycle checkpoints (4). In addition, genes that regulate DNA replication and chromosome segregation were also enriched in CIC-DUX4 replete cells compared to CIC-DUX4 knockdown (KD) cells (Figure 1B). In order to confirm these findings, we performed unbiased gene set enrichment analysis (GSEA) using 1426 up and down-regulated genes identified upon CIC-DUX4 KD in IB120 (endogenous CIC-DUX4) cells (Supplementary Table 1). This analysis consistently demonstrated enrichment of gene sets associated with G1/S, mitosis, cell cycle checkpoints, and DNA replication (Figure 1C). These expression data indicate that cell cycle and DNA replication programs are key molecular targets of the CIC-DUX4 fusion oncoprotein. Next, we queried a publicly available dataset that performed multiplex kinase activity (PamGene^TM^) profiling to identify active kinases in patient-derived CIC-DUX4 tumors, xenografts, and cell lines (22). Since our transcriptional analysis converged on genes involved in DNA replication, G1/S transition and mitosis, we focused on kinases that regulate the cell cycle in CIC- DUX4 sarcoma. Through a manual systematic analysis, we identified 20 unique phosphosites, mapping to 14 independent kinases previously implicated in cell cycle regulation (Figure 1D) (22). Multiple CDK1 (4) and CDK2 (3) phosphosites were identified in our analysis suggesting that these kinases have broad roles in CIC-DUX4 sarcoma cells. Interestingly, we noted that phosphosites mapping to the WEE1 kinase were highly repetitive and specific for the inhibitory phosphorylation site (Y15) on both CDK1 and CDK2 suggesting that CIC-DUX4 cells may depend on WEE1 activity to potentially delay G2/M progression, and limit premature mitotic entry and mitotic catastrophe (Figure 1E-F) (13, 15, 17). Consistent with this, we previously noted that ectopic expression of CIC- DUX4 leads to an increased G2/M fraction in NIH-3T3 cells (4). Since CIC-DUX4 transcriptionally upregulates *CCNE1,* which can induce a high replicative stress state in cancer (5, 6, 9–11,23), we hypothesized that WEE1 activity in CIC-DUX4 sarcomas may be an adaptive survival mechanism. To test whether WEE1 is necessary for CIC-DUX4 survival, we treated two independent patient-derived CIC-DUX4 cell lines (NCC_CDS1_X1_C1 and NCC_CDS2_C1) (22, 24) with the WEE1 inhibitor, adavosertib (AZD1775) (25–28). Adavosertib treatment significantly decreased viability of both NCC_CDS1_X1_C1 and NCC_CDS2_C1 cells as measured by Cell-Titer Glo (CTG) and crystal violet assays (Figure 2A-C). The decreased viability observed upon adavosertib treatment was associated with a decrease in CDK1 phosphorylation at Y15 and an increase in apoptosis as measured by PARP cleavage in CIC-DUX4-expressing NCC_CDS_X1_C1 and NCC_CDS2_C1 cells. Moreover, we observed an increase in the phosphorylation of Serine 139 on the histone variant H2AX (γH2AX), a sensitive marker of DNA damage, in adavosertib-treated cells compared to control (Figure 2D-E) (29). Coupled with enhanced Caspase 3/7 activity observed upon adavosertib treatment, these findings indicate that pharmacologic inhibition of WEE1 induces apoptosis in CIC-DUX4 sarcoma cells potentially through increased DNA damage and premature mitotic death (Figure 2F-G). To further mitigate possible off-target effects of adavosertib, we performed genetic silencing of WEE1 using two validated siRNAs (siWEE1-06, siWEE1-08) that target independent regions of WEE1 (30). Consistent with our pharmacologic studies, genetic inhibition of WEE1 decreased CDK1 Y15 phosphorylation and resulted in enhanced γH2AX expression and PARP cleavage compared to control (Figure 2H-I). Moreover, genetic silencing of WEE1 decreased NCC_CDS1_X1_C1 and NCC_CDS2_C1 cell viability compared to control as measured by Cell-Titer Glo assay (Figure 2J-K) and crystal violet (Figure 2L-M) assays. In order to further demonstrate that the adavosertib-mediated apoptotic effect was dependent on CDK1 and/or CDK2, we silenced CDK1 and/or CDK2 in CIC-DUX4-expressing cells and noted an increase in IC^50^ with combinatorial CDK1 and CDK2 silencing relative to control or either CDK1 or CDK2 KD alone (Figure 2N-O and Supplementary Figure 1A-C). In order to further link WEE1- mediated suppression of CDK1 activity to survival, we leveraged the well-characterized CDK1 variant, CDK1^AF^ (31). The WEE1 inhibitory phosphosite in CDK1 (Y15) is replaced in CDK1^AF^, generating a constitutively active isoform that is not responsive to WEE1 activity. Expression of this CDK1^AF^ variant in NCC_CDS1_X1_C1 or NCC_CDS2_C1 cells decreased viability compared to control (Figure 2P). These findings indicate that WEE1 may promote CIC-DUX4 survival through inhibitory phosphorylation of CDK1. Collectively, through these studies we reveal a pro-survival role for WEE1 and highlight a therapeutic vulnerability in CIC-DUX4 sarcomas.

**Figure 1.**
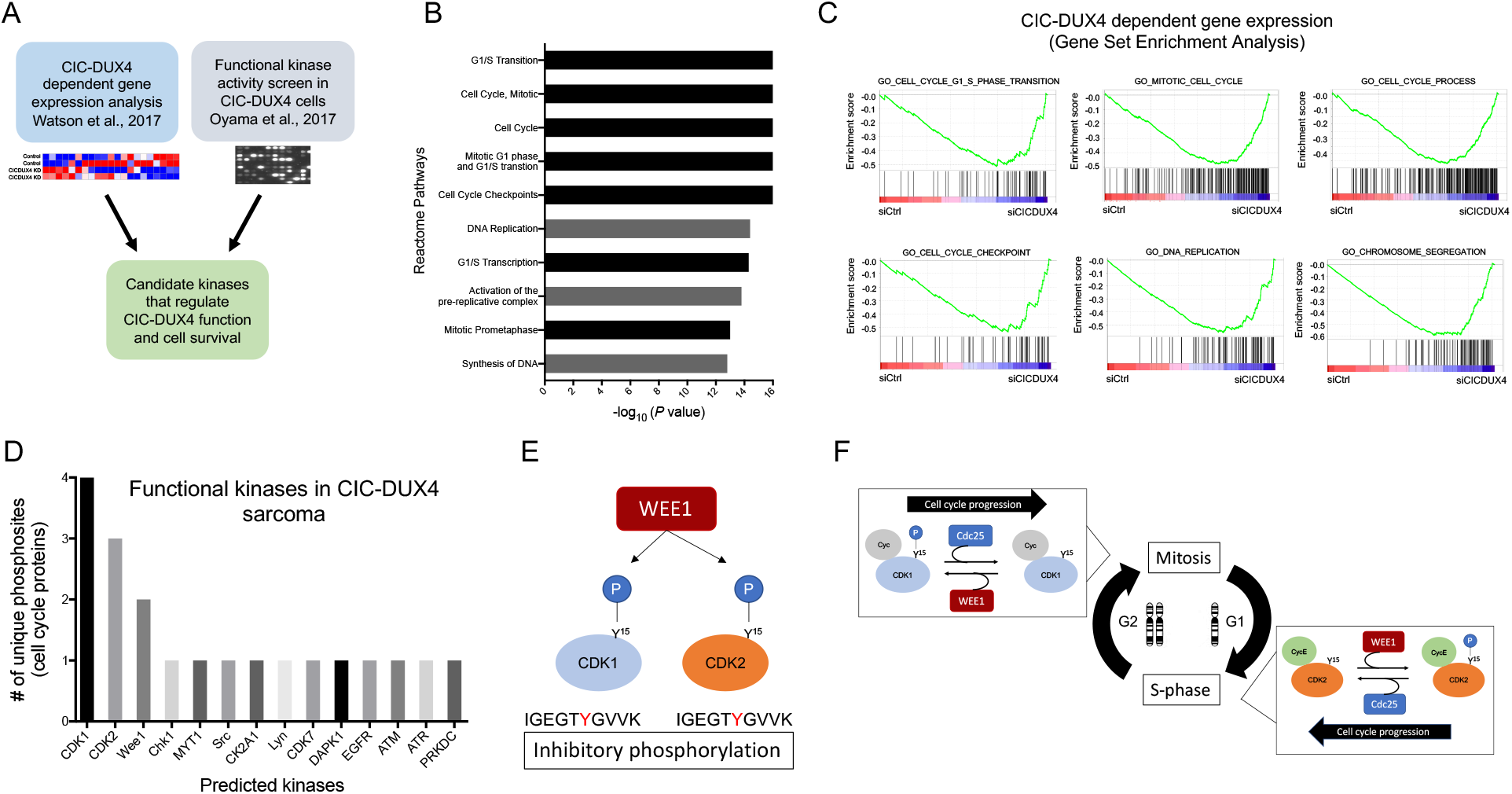
Integrative transcriptomic and kinase activity screen identifies Wee1 as a target in CIC-DUX4 sarcoma. (A) Approach to identify candidate kinases that regulate CIC-DUX4 survival. (B) Reactome pathway analysis identifies CIC-DUX4-dependent cell cycle and DNA replication pathways. (C) Gene Set Enrichment Analysis (GSEA) reveals cell cycle, DNA replication, and chromosome segregation at CIC-DUX4 targets. (D) PamGene array identifies functional kinases that regulate cell cycle and DNA replication in CIC-DUX4 cells. (E) Model of Wee1 inhibitory kinase motifs in CDK1 and CDK2. (F) Schematic of WEE1-regulated cell cycle control.

**Figure 2.**
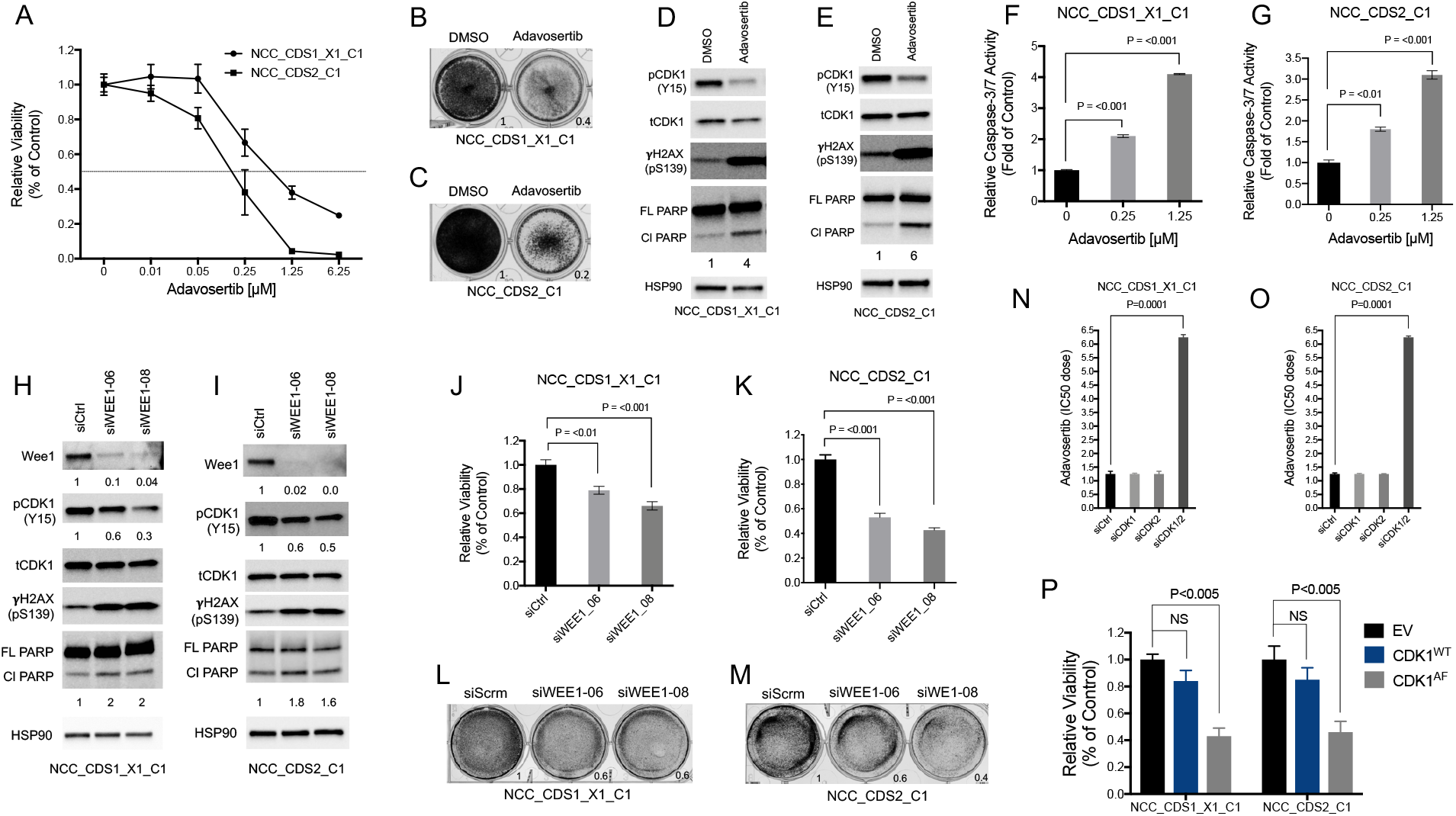
CIC-DUX4 sarcomas depend on WEE1 for survival. (A) Cell-Titer Glo viability assay of NCC_CDS1_X1_C1 and NCC_CDS2_C1 treated with adavosertib. Error Bars represent SEM. Performed in duplicate. Crystal Violet assay of NCC_CDS1_X1_C1 (B) and NCC_CDS2_C1 (C) cells comparing adavosertib (IC^50^ dose) and DMSO control. (D-E) Immunoblot analysis of NCC_CDS1_X1_C1 and NCC_CDS2_C1 cells treated with adavosertib or DMSO control. Representative figure; performed in duplicate. (F-G) Relative caspase 3/7 activity in NCC_CDS1_X1_C1 and NCC_CDS2_C1 treated with adavosertib vs control. P-value, 1-way ANOVA. Performed in triplicate (H-I) Immunoblot analysis of NCC_CDS1_X1_C1 and NCC_CDS2_C1 cells expressing two independent Wee1 siRNAs. Representative figure; performed in duplicate. (J-K) Cell-Titer Glo viability assay comparing two independent Wee1 siRNAs to scramble control in NCC_CDS1_X1_C1 and NCC_CDS2_C1 cells. P-value, 1-way ANOVA. Performed in triplicate. (L-M) Viability assay comparing two independent Wee1 siRNAs to scramble control in NCC_CDS1_X1_C1 and NCC_CDS2_C1 cells. P-value, 1-way ANOVA. Performed in duplicate. (N-O) Adavosertib IC^50^ dose in NCC_CDS1_X1_C1 and NCC_CDS2_C1 cells expressing siCDK1, siCDK2, or siCDK1 and siCDK2. P-value, 1-way ANOVA. Performed in triplicate. (P) Relative viability comparing NCC_CDS1_X1_C1 and NCC_CDS2_C1 cells expressing CDK1^WT^ or CDK1^AF^ versus vector control. P-value, 1-way ANOVA. Error bars represent SEM. Performed in triplicate.

In certain cancers, increased CCNE1 expression (amplification or transcriptional upregulation) deregulates the cell cycle at the G1/S and G2/M checkpoints by accelerating S-phase entry and stimulating premature mitosis (5, 7, 9–11, 23). Moreover, a hyperactive CCNE1/CDK2 complex can increase DNA origin firing-inducing rereplication, leading to high DNA replication stress – a state that requires cellular-based adaptive mechanisms to enable survival (32, 33). One response to this oncogene-induced replicative stress involves an increased dependence on the G2/M checkpoint kinase, WEE1 (15, 16). Since CIC-DUX4 directly binds to the regulatory region of *CCNE1* to hyperactivate the CCNE1/CDK2 complex (4), we hypothesized that CIC-DUX4-mediated *CCNE1* upregulation induces DNA replication stress, stimulating DNA repair responses and thus sensitizing to WEE1 inhibition. To mechanistically dissect the role of WEE1 in limiting extensive DNA damage and premature mitotic entry we first quantified γH2AX nuclear foci in NCC_CDS1_X1_C1 and NCC_CDS2_C1 cells through immunofluorescence (IF) staining following adavosertib treatment as previously described (11). We noted an increased fraction of cells with high nuclear γH2AX foci staining (>5 foci per cell) in adavosertib-treated cells compared to control (Figure 3A-B). These findings indicate that WEE1 activity can limit DNA damage in CIC-DUX4 sarcomas. Since WEE1 inhibition with adavosertib induces premature mitotic entry and DNA-damage (34, 35) in other cancer types, we next determined the impact of adavosertib on cell cycle progression in our patient-derived CIC-DUX4 sarcoma cells. Specifically, we analyzed the cell cycle distribution and γH2AX expression (DNA damage marker) in NCC_CDS1_X1_C1 and NCC_CDS2_C1 cells in response to adavosertib treatment. Compared to control, adavosertib (0.5μM at 48 hours) increased the G2/M fraction in our CIC-DUX4 cell lines (Figure 3C-E, I-K), thus indicating premature mitotic entry and/or mitotic arrest that was associated with extensive DNA damage (increased γH2AX expression) (Figures 3F-H, L-N).

**Figure 3.**
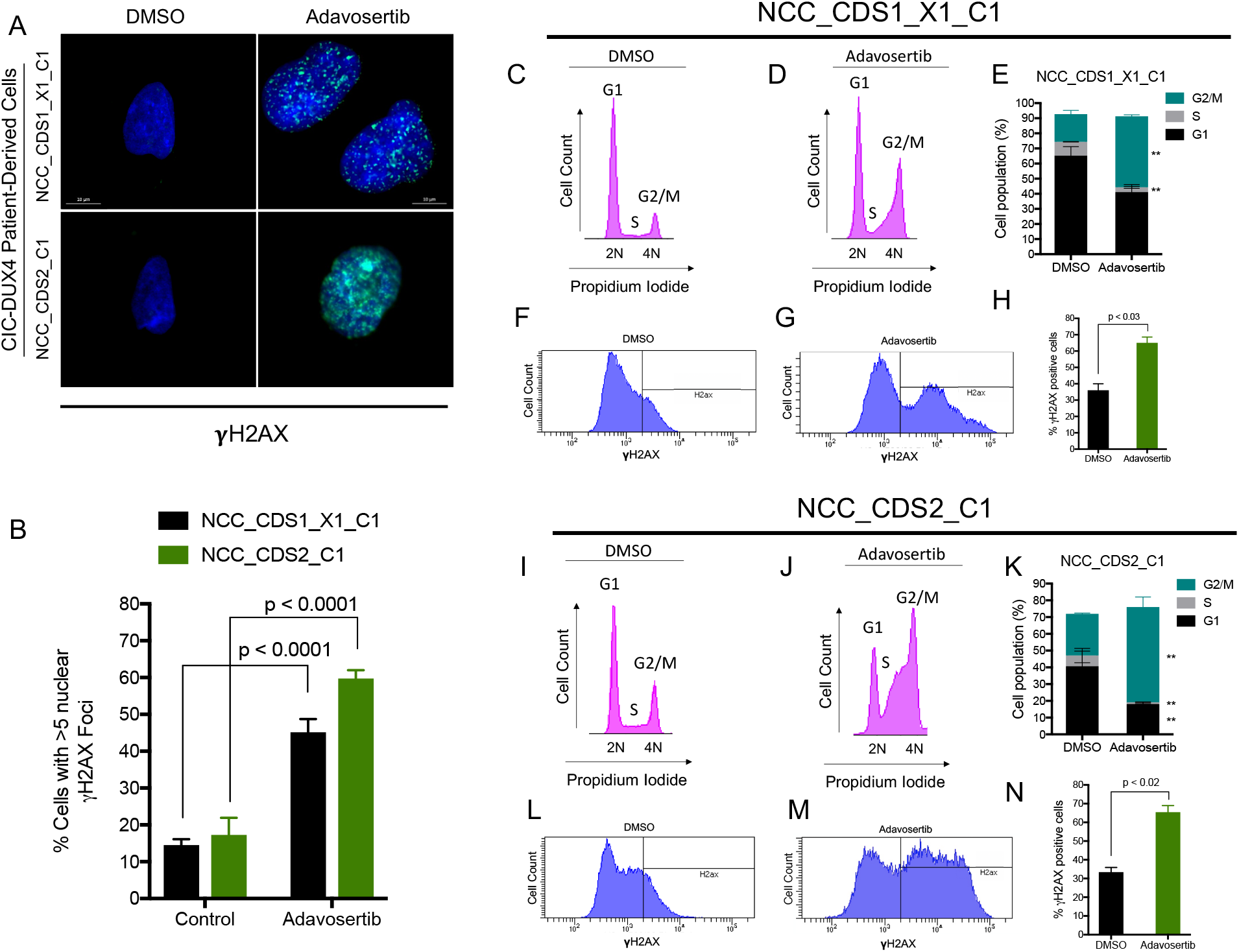
WEE1 inhibition increases DNA damage and mitotic entry in CIC-DUX4 sarcoma cells. (A) γH2AX immunofluorescence (IF) staining of NCC_CDS1_X1_C1 and NCC_CDS2_C1 cells treated with adavosertib (0.5μM) or DMSO (48 hours). Representative figure; performed in triplicate. (B) Quantitative analysis of γH2AX IF in (A) demonstrating the percentage of NCC_CDS1_X1_C1 or NCC_CDS2_C1 cells with >5 nuclear γH2AX foci following treatment with adavosertib or control. Mean percentage over 10 HPFs. P-value, student’s t-test. Representative cell cycle histograms (propidium iodide staining) of NCC_CDS1_X1_C1 (C-D) and NCC_CDS2_C1 (I-J) cells treated with adavosertib or DMSO (48 hours). Percentage of adavosertib (0.5μM) or DMSO-treated (48 hours) NCC_CDS1_X1_C1 (E) and NCC_CDS2_C1 (K) cells in G1, S, or G2/M phases as identified in (C-D and I-J). **P-value <0.05. Performed in duplicate. Error bars represent SEM. γH2AX expression in NCC_CDS1_X1_C1 (F-G) and NCC_CDS2_C1 (L- M) cells treated with adavosertib or DMSO. Percentage of γH2AX positive cells from NCC_CDS1_X1_C1 (H) and NCC_CDS2_C1 (N) cells analyzed in (F-G and L-M). P- value, student’s t-test. Error bars represent SEM.

CIC-DUX4 sarcomas are universally associated with poor clinical outcomes due to rapid metastatic progression and insensitivity to conventional chemotherapy agents (36). Thus, we next tested the translational impact of targeting the WEE1 kinase in preclinical models of CIC-DUX4 sarcoma. Specifically, we generated NCC_CDS1_X1_C1 and NCC_CDS2_C1 tumor xenografts in immunodeficient mice and treated with adavosertib (100mg/kg/day) and compared this to vehicle control. In NCC_CDS1_X1_C1 and NCC_CDS2_C1 bearing mice we noted tumor regressions in the adavosertib group compared to vehicle-treated mice (Figure 4A-B). The overall objective response rate (>/=30% reduction in tumor volume) to adavosertib in the NCC_CDS1_X1_C1 and NCC_CDS2_C1 cohorts were 87.5% (7/8) and 100% (6/6), respectively, without noted toxicity (Figure 4C-D and Supplementary Figure 2A-B). Consistent with our *in vitro* data, NCC_CDS1_X1_C1 and NCC_CDS2_C1 tumor explants from adavosertib-treated mice demonstrated a decrease in CDK1 phosphorylation (Y15) and increased PARP cleavage compared to vehicle control (Figure 4E-F). These *in vivo* studies further validate WEE1 as a therapeutic vulnerability in CIC-DUX4 sarcomas that can be readily targeted through clinically advanced WEE1 inhibitors, including adavosertib (28, 37). Collectively, these data demonstrate a mechanism-based therapeutic strategy to precisely and effectively target CIC-DUX4 sarcomas in patients.

**Figure 4.**
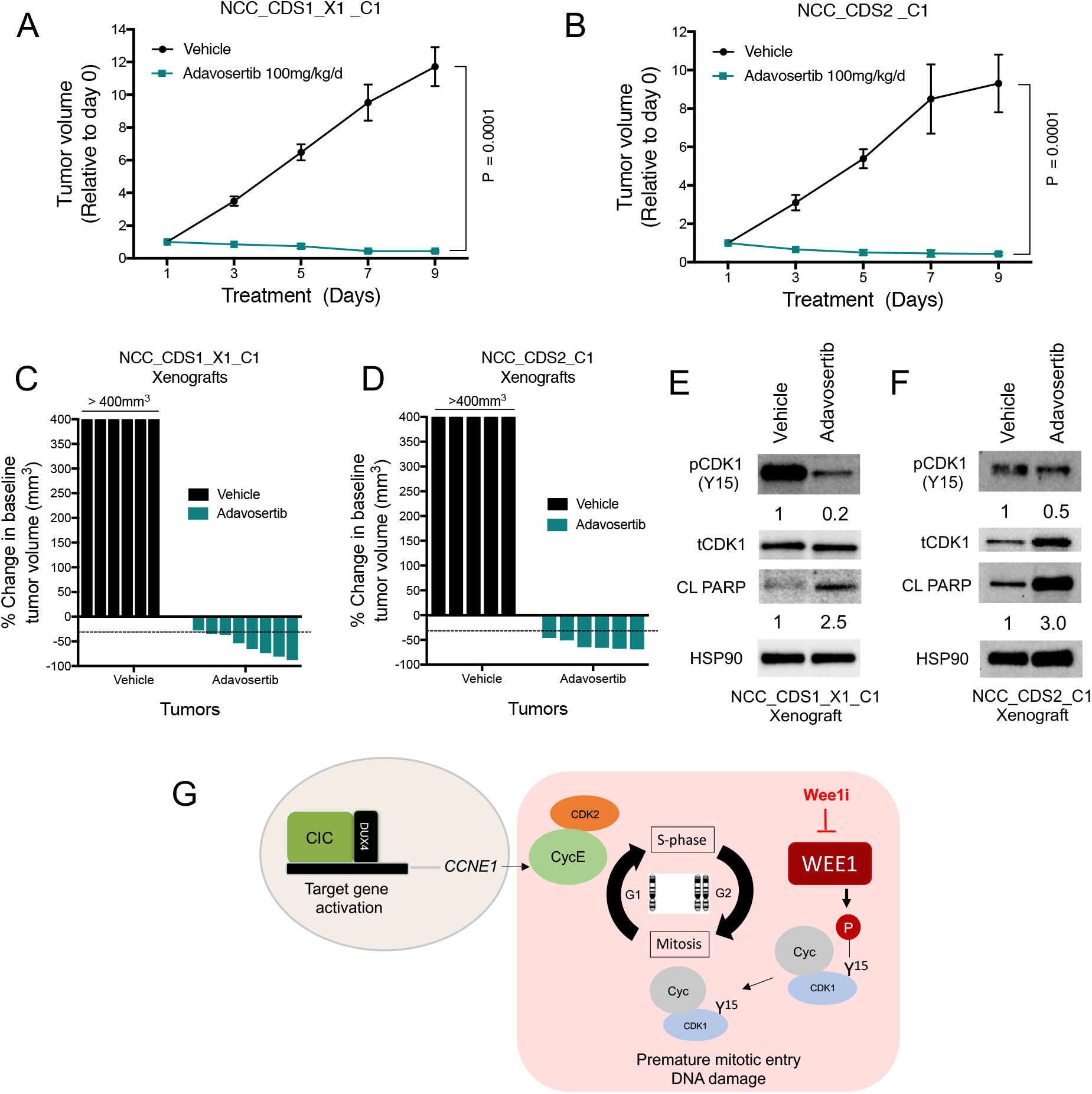
WEE1 is a therapeutic target in CIC-DUX4 sarcomas. Relative tumor volume of NCC_CDS1_X1_C1 (A) and NCC_CDS2_C1 (B) treated with adavosertib or vehicle control. P-value, student’s t-test, Error bars represent SEM. (C-D) Waterfall plots demonstrating the percent change in tumor volume for NCC_CDS_X1 and NCC_CDS2 tumor bearing mice comparing adavosertib and vehicle cohorts. (E-F) Immunoblot of NCC_CDS1_X1_C1 and NCC_CDS2_C1 tumor explants treated with adavosertib or Vehicle control. (G) Model of CIC-DUX4-regulated *CCNE1* transcriptional upregulation leads to a survival dependence on the G2/M checkpoint kinase, WEE1.

In this study, we employed an integrative transcriptional and functional approach to identify key clinically actionable vulnerabilities in CIC-DUX4 sarcoma. Through this analysis, we identified a WEE1-mediated adaptive response that enables CIC-DUX4 sarcoma survival by limiting massive DNA damage and mitotic catastrophe. These findings are in line with and expand our prior studies that reveal a dependence on CIC- DUX4-driven *CCNE1* expression in CIC-DUX4 sarcomas (4). Thus, we collectively demonstrate that CIC-DUX4 sarcomas transcriptionally upregulate *CCNE1*, compromising the G1/S checkpoint and conferring a dependence on WEE1 to limit DNA damage-associated cell death (Figure 4G). Importantly, these studies reveal a precision-based therapy for CIC-DUX4 sarcomas, which remain an aggressive and lethal subset of human cancer.

Through our studies we provide the initial translational framework for WEE1 inhibition as a therapeutic strategy in CIC-DUX4 sarcomas. Future studies should focus on rational combinatorial strategies to enhance DNA damage and potentially augment WEE1 inhibitor responses. Adavosertib is currently being evaluated in multiple tumor histologies and is proven safe as monotherapy or in combination with other conventional targeted and chemotherapy agents as well as radiotherapy (11, 27, 28, 37). Thus, adavosertib therapy is potentially a safe and efficacious therapy (28) that can be rapidly employed in the clinic to more effectively target CIC-DUX4 sarcomas. Similarly, ZN-c3 (Zentalis WEE1 inhibitor) is entering phase 2 studies as monotherapy and/or combination therapy in solid tumors, including sarcomas. One barrier in exploring the clinical response to WEE1 inhibitors in CIC-DUX4 patients is the relative rarity of CIC-DUX4 sarcomas (38). To overcome this obstacle, the sarcoma community must first differentiate CIC-DUX4 sarcomas as a unique entity and not routinely integrate (clinically or pathologically) them into more common small round cell sarcoma subtypes such as Ewing sarcoma (ES) (4, 18, 20, 36, 39). This misrepresentation leads to a misconception that CIC-DUX4 sarcomas should be managed and treated with a similar strategy as ES (36, 40, 41). In most cases, ES-directed chemotherapy is not effective in CIC-DUX4 sarcomas (36). Thus, in order to advance the field, we must develop a more rational approach to treat this ultra-rare yet lethal subtype. Additionally, extensive collaboration is warranted to more rapidly identify and direct CIC-DUX4 patients to clinical trials that may be effective such as WEE1 inhibitors (monotherapy or combination) or CDK2-directed therapies as previously described (4). Finally, perhaps a re-evaluation of clinical trial endpoints and a centralized clinical center for rare cancers would enhance clinical collaboration and accelerate therapeutic advancements (38).

## Methods

### Tumor xenografts

Six-to-eight week old female nude (NU/J) mice were purchased from Jackson Laboratory. Specific pathogen-free conditions and facilities were approved by the American Association for Accreditation of Laboratory Animal Care. Surgical procedures were reviewed and approved by the UCSF Institutional Animal Care and Use Committee (IACUC), protocol #AN178670-03B. For PDX subcutaneous xenotransplantation, 1.5×10^6^ NCC_CDS1_X1_C1 and NCC_CDS2_C1 cells were resuspended in 50% PBS/50% Matrigel matrix and injected into the flanks of immunodeficient mice.

### Cell lines and culture reagents

NCC_CDS1_X1_C1 and NCC_CDS2_C1 were generated as patient-derived cell lines by Tadashi Kondo (Oyama et al., 2017 and Yoshimatsu et al., 2020). The presence of the CIC-DUX4 fusion was confirmed in NCC_CDS1_X1_C1 and NCC_CDS2_C1 through RNAseq analysis using the “grep” command as previously described (Panagopoulos et al, 2014). NCC_CDS1_X1_C1 cells were maintained at 37 °C in a humidified atmosphere at 5% CO2 and grown RPMI 1640 media supplemented with 10% FBS, 100 IU/ml penicillin and 100ug/ml streptomycin. NCC_CDS2_C1 cells were cultured in RPMI media supplemented with 10% FBS, 100 IU/ml penicillin and 100ug/ml streptomycin. Adavosertib (AZD1775, MK1775) was purchased from MedChemExpress (HY-10993).

### Gene knockdown and over-expression assays

ON-TARGET scramble, WEE1 (WEE11-06-J-005050-06 and WEE1-08-J005050-08), CDK2 (L-003236-00-0005), and CDK1 (L-003224-00-0005) siRNAs were obtained from GE Dharmacon and transfections were performed with Lipofectamine RNAiMAX transfection reagent. CDK1 (Cat. 61840) and CDK1^AF^ (Cat. 39872) plasmids were purchased from Addgene and transfections were performed with FuGENE 6 transfection reagent.

### Western blot and qRT-PCR

All immunoblots represent at least two independent experiments. Adherent cells were washed and lysed with RIPA buffer supplemented with proteinase and phosphatase inhibitors. Proteins were separated by SDS-PAGE, transferred to Nitrocellulose membranes, and blotted with Cell Signaling Technology (CST) antibodies recognizing: HSP90 (CST #4874), total CDK1 (CST #9116), phosphorylated (Y15) CDK1 (CST #9111), WEE1 (CST #13084), PARP (CST #9542), total H2AX (CST #7631), and phosphorylated S139 γH2AX (CST #9718).

### Xenograft tumors

Subcutaneous xenografts were explanted at the study end-point and immediately immersed in liquid nitrogen and stored at −80 degrees. Tumors were disrupted with a mortar and pestle, followed by sonication in RIPA buffer supplemented with proteinase and phosphatase inhibitors. Proteins were separated as above. Antibodies to PARP, phosphorylated (Y15) CDK1, HSP90, Total CDK1 were purchased from Cell Signaling Technology.

### Cell viability assays

Cells were seeded overnight at a density of 3,000 cells per well in 96-well plates and treated with relevant agents for 72 hours. Cell viability was determined using the CellTiter-GLO (Promega) assay according to the manufactures protocol. Each assay consisted of at least three replicate wells. For crystal violet assays, 100,000 cells were seeded per well in a 12-well plate (250,000 cells in a 6 well plate) and allowed to grow for five consecutive days. Cells were then fixed in 3.7% paraformaldehyde, followed by 0.05% Crystal Violet stain. Quantification was performed using Image J software.

### Apoptosis assays

Cells were seeded overnight at a density of 40,000 cells per well in 96-well plates and treated with relevant agents for 24 hours. Caspase 3/7 activity was measured on a Molecular Devices microplate reader using Caspase Glo reagent from Promega per the manufactures protocol and normalized to cell number.

### Immunofluorescence

Immunofluorescence was performed on glass coverslips. Cells were fixed with 4% paraformaldehyde, quenched with 1X PBS and 10mM glycine and permeabilized with 0.1% Triton X-100, then incubated with conjugated (Alexa Fluor 488) phospho Histone H2A.X (S139) antibody (CST). ProLong Gold Antifade Mountant with DAPI was applied directly to fluorescently labeled cells on microscope slides. Fluorescent images were collected on Zeiss Axioplan II fluorescent microscope.

### Cell cycle analysis

To determine the effect of adavosertib (0.5 μm for 48 hours) on the cell-cycle of NCC_CDS1_X1_C1 and NCC_CDS2_C1 cells, we first trypsinized, washed with PBS +0.1% BSA and fixed in ice cold ethanol overnight. We subsequently treated with RNAse (Cell Signaling) and stained with propidium iodide (PI) solution (Thermo Fisher Scientific) or stained with conjugated (Alexa Fluor 488) phospho Histone H2A.X (S139) antibody (CST) at room temperature for 30 minutes. Cells were analyzed on a BD LSRII flow cytometer.

### Pathway and Gene Set Expression Analysis (GSEA)

As previously described in Okimoto et al., 2019, 165 down-regulated genes were identified in IB120 cells expressing siCtrl or siCIC-DUX4. 43/165 putative gene targets contained the CIC DNA-binding motif (T[G/C]AATG[A/G]A) within −2 kb and +150 bp of the transcription start site. These 43 high confident genes were analyzed using Reactome Pathway Software to identify CIC-DUX4 regulated pathways. Gene set enrichment analysis (GSEA) was performed as previously described (Subramanian et al., PNAS 2005) using the top 1426 up-and-down regulated genes in IB120 cells expressing siCIC- DUX4 vs control.

### Kinase array analysis

As previously described in Oyama et al., 2017, multiplex kinase activity profiling was performed in CIC-DUX4 sarcoma patient-derived cells using PamGene^TM^ technology and measured by PamStation according to the manufactures protocol. Manual identification of cell cycle kinases through the phosphorylation status of substrate peptides was performed using the published Oyama dataset.

### Statistical analysis

Experimental data are presented as mean +/- SEM. P-values derived for all *in-vitro* and *in vivo* experiments were calculated using two-tailed student’s t-test or one-way ANOVA.

## Author contributions

R.K.P and N.J.T. designed and performed the experiments and analyzed the data. NQB analyzed the data and provided critical revisions on the manuscript. T.K. performed experiments and provided cell lines. R.A.O directed the project, analyzed experiments, and wrote the manuscript.

## Acknowledgements

R.A.O. was supported by grants from the National Cancer Institute K08CA222625. N.J.T was supported by a Research Fellowship from the UCSF School of Medicine.

