## Supplemental Figures for "WEE1 kinase is a therapeutic vulnerability in CIC-DUX4 undifferentiated sarcoma"

The Supplemental material contains two figures and one table.

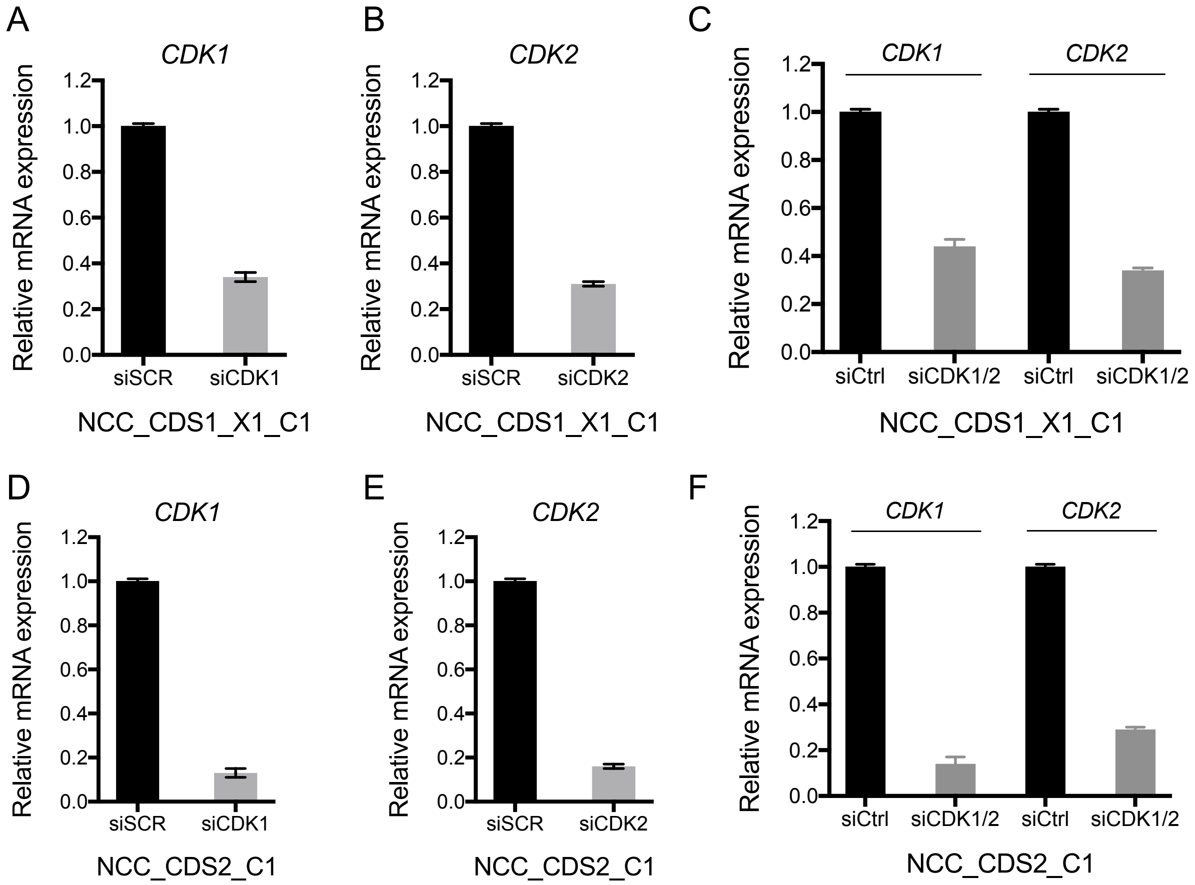


**Supplementary Figure 1. CDK1 and CDK2 genetic silencing in CIC-DUX4 cells.**

Relative mRNA expression of CDK1 (A), CDK2 (B), and CDK1 and CDK2 (C) in NCC_CDS1_X1_C1 cells following siRNA-mediated knockdown. Relative mRNA expression of CDK1 (D), CDK2 (E), and CDK1 and CDK2 (F) in NCC_CDS2_C1 cells following siRNA-mediated knockdown.

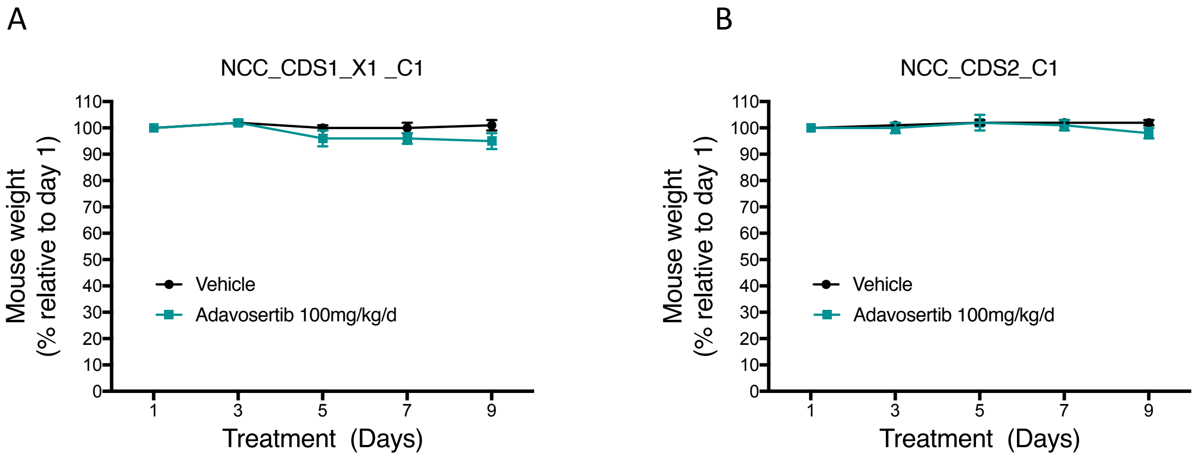


**Supplementary Figure 2. Adavosertib is well tolerated in mice.**

Serial weights of mice harboring NCC_CDS1_X1_C1 (A) and NCC_CDS2_C1 (B) xenografts treated with adavosertib or vehicle control.

**Supplementary Table 1. 1426 differentially expressed genes in IB120 cells +/- CIC-DUX4 KD.**
